# Harnessing Protein Unfolding for Thermosensing: Structural Insights from TRPV3

**DOI:** 10.1101/2025.09.17.676191

**Authors:** Andrew Njagi Mugo, Dinesh Indurthi, Ryan Chou, Gaya P. Yadav, Liguo Wang, Feng Qin

## Abstract

How proteins sense temperature with high precision is a fundamental biological challenge. Here, we structurally elucidate a dynamics-based mechanism for thermoTRPs, termed suicidal gating, in which extreme thermosensitivity arises from intrinsic instability and concerted protein dynamics, coupling channel opening to partial protein unfolding. Using cryo-EM, we directly capture heat-induced partial unfolding within the channel’s central pore domain, a structural core of gating. Analysis of mutant structures capturing intermediate activated states reveals no large domain shifts despite high temperature dependence, only subtle interfacial changes between domains. Both mutation and heat treatment consistently impact a critical network of thermolabile "latch" interactions along the S6-TRP helix-proximal N-terminus axis, driving receptor activation from a stable closed structure to progressively decoupled states and eventual disintegration. These findings provide direct structural evidence for a new receptor mechanism and establish a broader paradigm in which distributed structural flexibility, rather than localized dedicated sensors, drives extreme biological functions.

## Introduction

Temperature receptors serve as the body’s frontline detectors of thermal and noxious stimuli. Most identified receptors to date belong to the transient receptor potential (TRP) superfamily, or thermoTRPs ^1,2^. Heat-sensitive vanilloid channels (TRPV1-3) detect warmth and noxious heat ^3-7^, while melastatin channels (TRPM) include both cold-sensitive members such as TRPM8 ^8,9^ and heat-sensitive channels like TRPM4 ^10-15^. Each channel is tuned to a specific temperature range, collectively allowing the thermoTRP family to cover the full spectrum of physiologically relevant temperatures. Beyond acute sensing, many thermoTRPs also mediate nociceptive signaling in pathological states ^16^, positioning them as prime targets for innovative analgesic therapies ^17^.

Despite their central role in thermosensation, the molecular basis of temperature detection by thermoTRPs remains unresolved. Central to this challenge is their extraordinarily steep temperature dependence, which thermodynamically requires activation energies far beyond ordinary biochemical processes ^18,19^. This extreme sensitivity has inspired sensor-centric models that posits specialized thermosensory domains—akin to voltage-sensing domains in voltage-gated channels—to detect temperature ^20,21^. The heat capacity model, in particular, attributes gating to energetic changes arising from the burial/exposure of nonpolar residues ^22-25^. Yet, decades of mutagenesis and structural studies implicate residues broadly distributed throughout the channel ^26-44^. Consequently, no consensus thermosensory domain has emerged, and a unifying mechanism remains elusive.

Cryo-electron microscopy (cryo-EM) offers unprecedented access to the structural underpinnings of temperature sensing. TRPV1 studies reveal channel opening upon combined agonist and heat activation ^39^, while TRPV3 structures capture heterogeneous, heat-induced states resembling functional intermediates ^37,38^. However, structural rearrangements across thermoTRPs remain variable, and their contributions to extreme temperature sensitivity remain unclear. Moreover, the pronounced time-dependence of thermal activation, including fast gating ^19,45^, sensitization ^46-48^, and unfolding ^49-52^, creates a temporal mismatch with the equilibrium or metastable states captured by cryo-EM, complicating mechanistic interpretation.

Recent calorimetry shows that thermoTRP channels are intrinsically thermally unstable, undergoing partial unfolding within their physiological detection range ^49,50^. This instability underlies a ‘suicidal’ gating mechanism, in which channel opening is coupled to thermal unfolding, accounting for the characteristic functional irreversibility and the large enthalpic changes that drive extreme temperature sensitivity. Unlike voltage- or ligand-gated channels that rely on discrete sensors, temperature-dependent gating harnesses global, concerted protein dynamics, consistent with broadly distributed thermosensitive residues revealed by mutagenesis. Evidence that TRPM8 cold sensing also operates through dynamic fluctuations further highlights the generality of this mechanism across thermoTRP channels ^53^.

Here, we interrogate the structural basis of suicidal gating, revealing how channel activation is coupled to partial protein unfolding, where unfolding occurs, and whether large-scale conformational changes are required for extreme temperature sensitivity. To overcome cryo-EM limitations in capturing transient states, we combine structural analyses with engineered mutant channels that capture activated intermediate states, enabling a direct link between structure and function. We show that localized interfacial dynamics, particularly along the pore–TRP helix–proximal N-terminus axis, drive high temperature dependence and identify specific thermolabile latches orchestrating temperature-dependent state transitions. These findings provide the first structural blueprint for suicidal gating and establish a distinct paradigm in which global structural instability, rather than localized dedicated sensors, governs extreme biological function.

## Results

### Structural mapping of heat-induced partial unfolding

We first asked whether partial unfolding occurs during heat activation, as predicted by DSC, and sought to pinpoint where it takes place within channels. Wild-type channels undergo highly concerted, domain-spanning thermal transitions at energies consistent with extensive unfolding ^49,50^, which can obscure localized structural changes and require high activation temperatures prone to nonspecific effects. To circumvent these limitations, we targeted the membrane-proximal N-terminus, a key coupling hub ^26,28,49^, using the TRPV3-412S mutant containing a single-residue insertion ^27^. This minimal perturbation preserved heat activation while reducing temperature sensitivity, effectively attenuating interdomain coupling and lowering the activation threshold, enabling us to limit global unfolding and directly visualize discrete structural transitions within the channel.

We purified and reconstituted the TRPV3-412S mutant into nanodiscs, and heated the sample to 42 °C, a temperature selected within the channel’s functional activation range ^27^. Heating was performed via differential scanning calorimetry at 1 °C/min ^50^ to promote population of diverse kinetic states, followed by plunge-freezing for cryo-EM imaging. Single-particle analysis identified two distinct structural classes; here, we focus on the dominant class, with the minority class described below.

Fig. 1a-d show the reconstructed density maps of the channels. Compared with the wild-type resting state (Fig. 1a,c), the heated sample revealed a striking absence of density in the central pore (Fig. 1b,d), particularly within the lower cavity (Fig. 1b). We interpreted this loss of signal as indicative of elevated mobility, reflecting tertiary structural collapse or partial unfolding. In contrast, densities for the peripheral S1-S4 transmembrane bundle and intracellular domains – including the ankyrin repeat domain (ARD), proximal N-terminus, and C-terminus – remained robust, highlighting the pore’s selective susceptibility to thermal destabilization.

**Figure 1.**
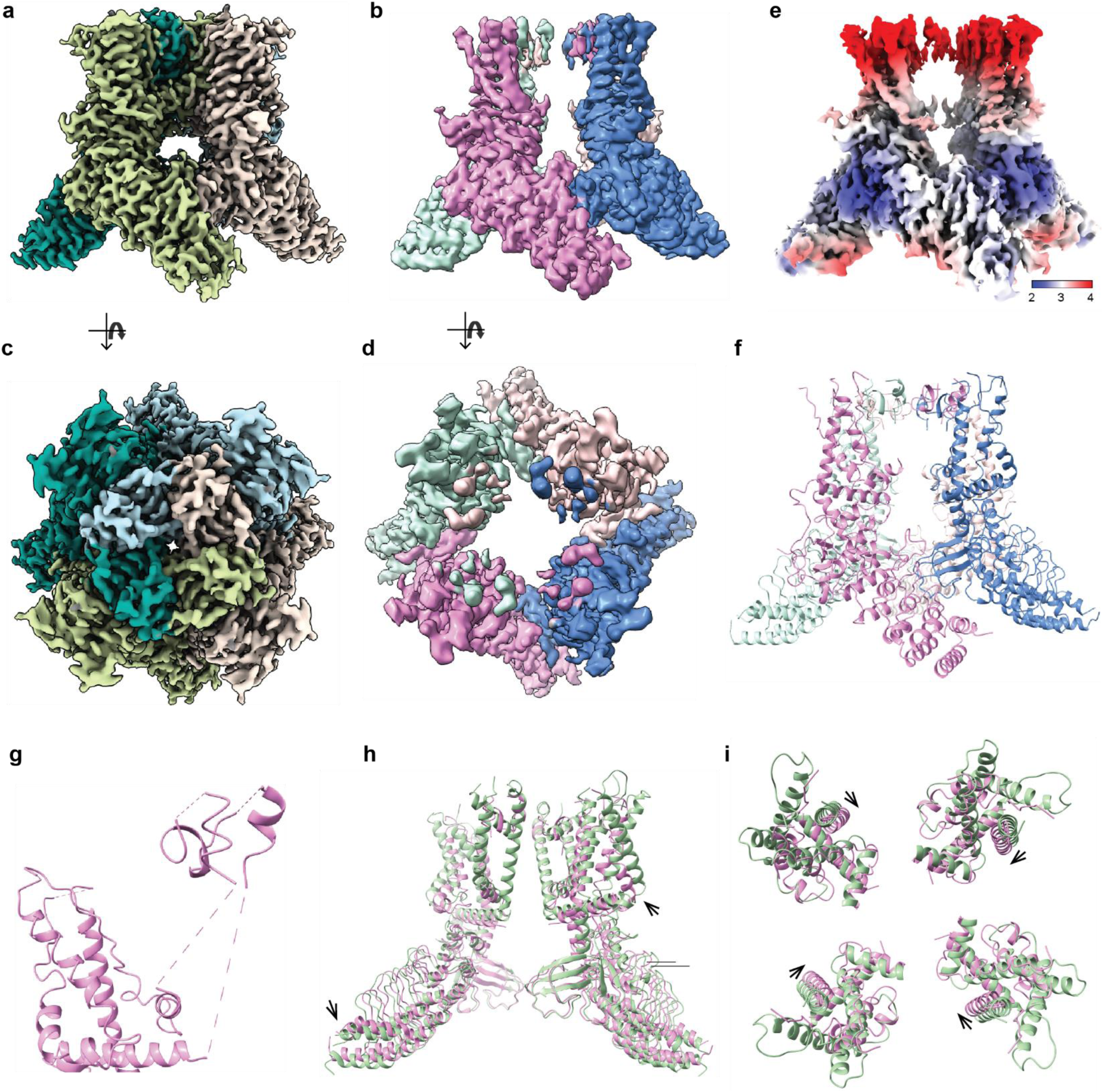
Partial unfolding of TRPV3 channels upon heating. **a-d)** Cryo-EM density maps of wild-type (WT) TRPV3 in resting state (a) and TRPV3-412S mutant after slow heating to 42 °C (1 °C/min, DSC) (b), showing substantial loss of density in the central pore domain. Subunits are individually color-coded for clarity; maps shown in side view parallel to membrane (a,b) and in top view from extracellular side (c,d). **e)** Local resolution map of TRPV3-412S. **f)** Atomic model of DSC-heated (denatured) state in side view, rendered as a ribbon diagram. **g)** Close-up of the pore region, highlighting fragmented S5 (residues 595-601) and S6 (residues 661-686) helices, as well as distorted S4 with unresolved side chains. **h,i)** Superimposition of DSC-heated TRPV3-412S (pink) with WT (green) in side (h) and bottom (intracellular) (i) views. Arrows indicate global changes upon heating, including overall vertical shortening (h) and lateral contraction of the transmembrane domain (i).

Fig. 1e depicts the local resolution map in the heated structure, revealing marked heterogeneity, with distal transmembrane helices (especially S3 and S4) resolved at a lower resolution of ∼3–4 Å versus intracellular domains at ∼2–3 Å. Nevertheless, most domains were traceable for molecular modeling (Fig. 1f). The model indicated that unfolding began after the S4-S5 linker (residue 584) and extended through S5, S6, and into the N-terminal start of the TRP helix (residue 688) (Fig. 1g). The upper extracellular pore retained partial density at low resolution, suggesting that S5 and S6 were the most destabilized, whereas the S4-S5 linker, proximal TRP helix, and pore helix rim remain relatively more stable and act as structural barriers to confine unfolding to the pore core. Furthermore, while S5/S6 density was absent at standard contour levels, lowering the threshold revealed diffuse density connecting the S4-S5 linker/TRP helix to the upper pore (Fig. S1-a, boxed). Modeling this density with poly-alanine allowed backbone tracing consistent with S5 and S6 positions in resting-state structures (Fig. S1-b, red), confirming that these helices persisted but were only highly mobile.

Outside the pore, the S1-S4 bundle was largely intact but displayed substantial distortion in S3 and S4 (Fig. 1f,g), likely from weakened or lost pore contacts. Other regions – including S1-S2, the ARD, and the TRP helix-MPD complex – were well-resolved and maintained resting-state conformations.

Heating also produced subtle global architectural changes (Fig. 1h,i). Superimposing heated TRPV3-412S (pink) onto resting wild-type (green) revealed mild vertical compression (Fig. 1h, arrows) and lateral contraction toward the pore axis (Fig. 1i, arrows), mirroring changes reported for heated wild-type TRPV3 and TRPV1 ^37-39^. These alterations likely reflect thermal relaxation to a lower-energy state – essentially an annealing effect. Overall, at functional activation temperatures, the channel exhibits domain-specific unfolding that originates in the pore (S5/S6) and progresses toward peripheral helices, accompanied by modest global compaction. As the pore is central to gating, these changes provide direct structural basis linking activation to unfolding, supporting the suicidal gating mechanism.

### High temperature dependence without major domain shifts

The suicidal unfolding model posits that extreme temperature sensitivity arises from coordinated, distributed protein dynamics rather than large-scale movements of discrete sensor domains. Directly capturing these transient, heat-activated states by cryo-EM is challenging, as prolonged heating risks nonspecific changes and denaturation. To overcome this, we exploited TRPV3 sensitization, in which heating drives a resting-to-sensitized transition that underlies high temperature dependence ^27^. The TRPV3-412S mutant functionally mimics this sensitized state without applied heat, providing a stable structural surrogate to directly capture thermosensitive conformations while avoiding complications inherent to wild-type channels.

Using this approach, we determined cryo-EM structures of nanodisc-reconstituted WT TRPV3 (Fig. 1a,c) and TRPV3-412S (Fig. 2a) under resting (non-heated) conditions, enabling direct comparison of structure features relevant to thermosensitivity. Both structures exhibited well-resolved density for residues 117-758, excluding distal termini and select extracellular loops, with overall resolutions of 2.9 A (WT) and 3.0 A (mutant) (Fig. 2b, Fig. S2-a). The structures closely resembled each other (RMSD=1.02Å) and prior TRPV3 models ^37,38,54-61^. In both, the selectivity filter (SF) was dilated at Gly638 (WT) or Gly639 (412S) with a radius of ∼7 - 8 Å, consistent with an open upper gate, while the lower gate at Met677 (WT) / Met678 (mutant) was constricted (∼5.0 Å WT, ∼6.0 Å mutant) (Fig. 2c,d, Fig. S3), indicating a non-conducting state, albeit slightly wider in the mutant. The S6 helix maintained an α-helical form in all structures.

**Figure 2.**
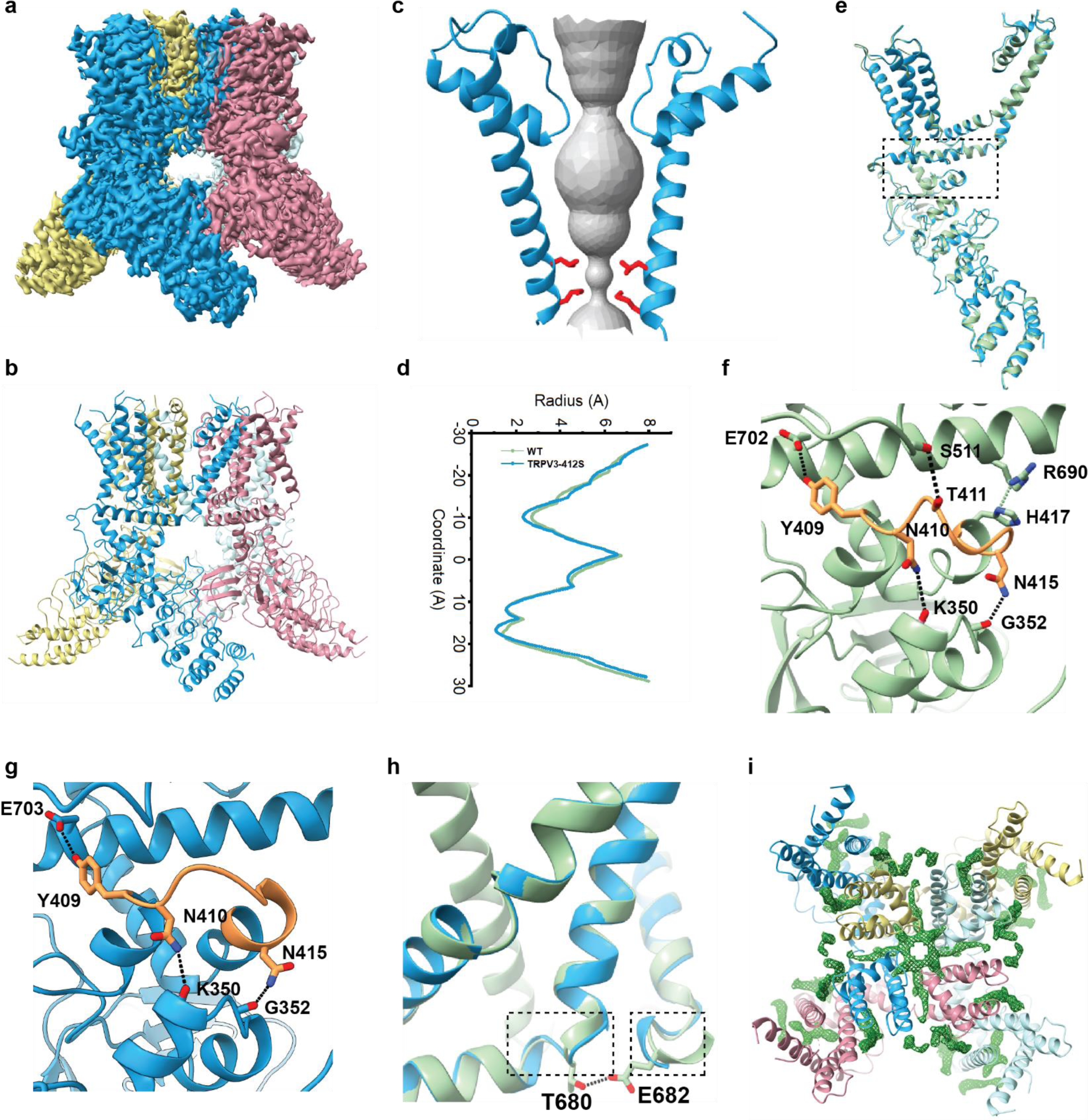
Structural basis of high temperature dependence. **a-b)** Cryo-EM density map of TRPV3 mutant capturing a heat-activated, temperature-dependent intermediate seen in wild-type (a) and cartoon representation of its atomic model (b). Subunits are individually color-coded. **c,d)** Space-filling representation of the pore (c) and corresponding pore radius profile along the channel axis (d, blue) superimposed on the wild-type distribution (green). **e)** Superimposition of the mutant structure (blue) with the WT resting state (green) within a single protomer, revealing no major domain rearrangements but localized differences at key domain interfaces (boxed), including proximal N-terminus contacting TRP helix and S2-S3 loop, and S6-TRP elbow linker. **f,g)** Detailed views of interfacial “latches”. In WT (f), the proximal N-terminus engages the TRP helix at both ends (Tyr409—Glu702 and His417—Arg690) and the S2-S3 loop (Thr411—Ser511). In the mutant (g), only Tyr409—Glu702 persists. The S6-TRP elbow linker is also altered (h, boxed), showing slight helix unwinding/rewinding at the junction. **i)** Lipid-like densities (green) in the pore region, S1-S4 bundle, and vanilloid binding cavity, shown in top/extracellular view.

Superimposition (Fig. 2e) revealed subtle, but position-critical differences at the interfaces (boxed) involving membrane-proximal N-terminus (MPD), TRP helix, and adjacent TMD elements. In WT (Fig. 2f), a helix-loop-helix (HLH) motif (residues 408–417) in the MPD formed a stabilizing network with the TRP helix (His417–Arg690, Tyr 409–Glu702), S2-S3 loop (Thr411–Ser511), and ARD (Asn410-Lys350, Asn415-Gly352), anchoring proximal N-terminus at a key gating interface.

In TRPV3-412S (Fig. 2g), the HLH motif was remodeled: the C-terminal helix elongated and the inter-helical loop shortened, loosening or disrupting the stabilizing interactions observed in WT. The mutant HLH adopted three conformations (Fig. S4): a predominant state retaining partial contact with TRP helix and ARD; a further disengaged state retaining a single ARD contact; and a more extended, solvent-exposed state with similar interactions. Even in the predominant state (Fig. 2g), key interactions with S2-S3 loop and N-terminus of TRP helix were lost. This increased flexibility propagated to the TRP helix N-terminus, inducing partial unwinding at the S6 junction and slight elongation of the TRP helix N-terminus (Fig. 2h), effectively shifting the S6-TRP loop upward towards the gate. Notably, these HLH-centered structural changes align with prior functional data implicating this motif in strong temperature dependence ^26^, supporting a role for its local interfacial latching in thermal gating.

Both WT and mutant structures featured non-proteinaceous densities consistent with annular lipids in transmembrane crevices (WT: Fig. S2; mutant: Fig. 2i, Fig. S5); Due to limited resolution, lipid head and tail groups could not be unambiguously assigned and were not modeled). Two prominent lipid-like densities matched previously reported sites: one between the extracellular half of S4 and the pore of a neighboring subunit, and another between the intracellular S1-S4 domain and the TRP domain ^37,38,54,56,57^. Additionally, we observed a novel ring-shaped density, presumably lipids, positioned laterally beneath the pore helix with “finger-like” projections extending into intersubunit spaces between S6 and adjacent S5/S6 (Fig. 2i; Fig. S2-b,d). This arrangement resembles lateral fenestrations in potassium channels ^62^, calcium uniporter ^63^, sodium channels ^64^, and TRPM8 ^65^, placing these lipids near gating-relevant regions and suggesting possible roles in lipid- or drug-mediated modulation. Interactions with S6 may contribute to stabilizing the observed α-helical conformation in all structures.

Despite their potential functional relevance, these lipid-like densities were conserved across WT and mutant structures, consistent with prior functional evidence that temperature dependence is intrinsic and independent of membrane fluidity or native lipid composition ^66,67^. Together, the close structural similarity, localized interfacial remodeling at the MPD–TRP–S6 junction, and preserved lipid environment are consistent with a model in which temperature dependence is encoded primarily through protein dynamics rather than by major domain rearrangements or lipid composition. In the mutant, weakened interdomain coupling likely lowers the energetic cost of activation, thereby facilitating gate opening and mimicking the sensitized WT state.

### Heat-induced interfacial remodeling

To capture heat-induced conformations beyond those stabilized by mutation, we directly activated nanodisc-reconstituted TRPV3-S412 channels by heating prior to plunge-freezing. We used two approaches: slow ramping via DSC (1 °C/min) to populate diverse kinetic states, and rapid heating by pre-equilibrating the Vitrobot chamber, both targeting 42 °C. Because thermal activation tends to be irreversible, cryo-EM analysis of these samples was expected to reveal gating-relevant conformations underrepresented in the sensitization-mimicking mutant.

DSC heating yielded predominantly denatured particles (Fig. 1b,d), but a minor fraction produced a well-resolved, intact reconstruction. Vitrobot heating similarly yielded an intact class. From these datasets, we solved two high-resolution structures at 2.5 Å: intermediate state 1 (IM1, from Vitrobot heating; Fig. 3a) and intermediate state 2 (IM2, from DSC heating; Fig. 3c), designated based on their extent of structural changes.

**Figure 3.**
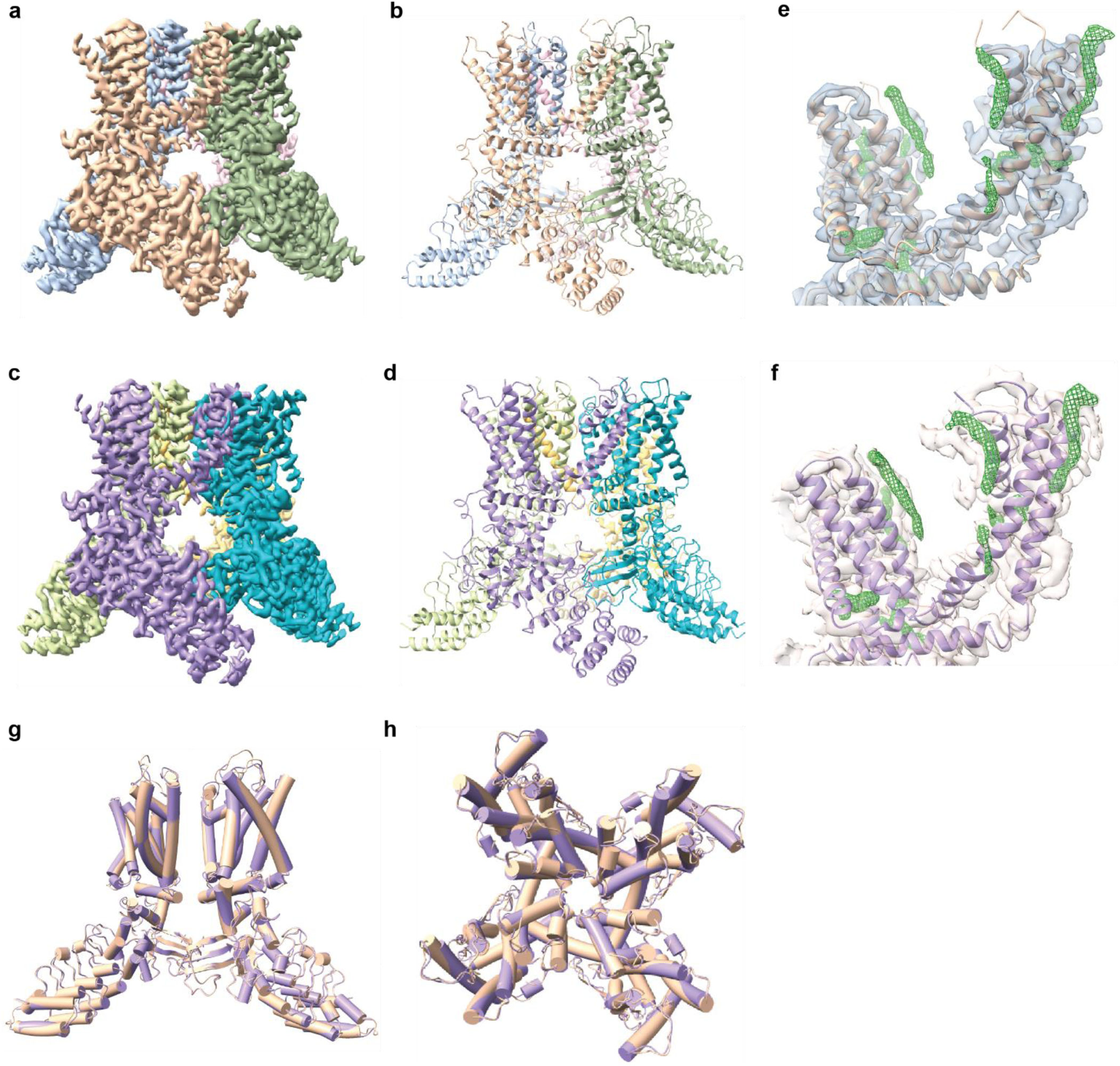
Heat-activated intermediate states without large-scale domain shifts. **a-d)** 3D cryo-EM reconstructions of TRPV3-412S after heat activation by two distinct protocols: rapid heating in a Vitrobot chamber (a) and slow heating by differential scanning calorimetry (DSC, 1 °C/min) (c), with corresponding atomic models fitted to the maps (b,d). Subunits are individually color-coded. **e-f)** Lipid-like densities (green) observed in both heat-activated structures (Vitrobot, e; DSC, f), located around the pore domain, S1-S4 bundle, and vanilloid binding cavity between S2 and S3. **g-h)** Superimposition of the two heat-activated structures (Vitrobot, tan; DSC, purple), showing nearly identical overlap, indicating that both heating protocols stabilize a common intermediate. The overall architecture of these heat-activated states also closely resembles that of unheated channels, with no large-scale structural rearrangements.

Both heated structures (IM1, Fig. 3b; IM2, Fig. 3d) closely resembled the resting mutant (RMSD: IM1=1.96 Å; IM2=2.20 Å) and retained constricted lower gates (Fig. S6), confirming a non-conducting state. In IM2, the narrowest constriction at Met678 approached that of Leu674 (Fig. S6-d), resulting a slightly wider aperture, likely reflecting increased flexibility at the S6-TRP helix junction (elaborated below). Lipid-like densities (Fig. 3e,f; Fig. S7), including the ring-shaped features beneath the pore helix, were preserved, suggesting no gross lipid rearrangements. Notably, despite markedly different heating rates, IM1 and IM2 were nearly identical (RMSD = 0.96 Å) (Fig. 3g, h), converging on a common structure, suggesting that the channel involves only a narrow set of thermally accessible metastable states.

The differences between heated and nonheated structures, as well as between IM1 and IM2, were confined to key gating interfaces also affected by the S412 mutation (Fig. 4). At the S6-TRP helix junction (Fig. 4a), IM1 exhibited partial S6 unwinding from the helix bundle crossing, further elongating the S6– TRP elbow loop. Concurrently, the S2-S3 loop displayed a tendency toward helix formation (Fig. 4a), and in the proximal N terminus (Fig. 4b), the HLH motif unwound along its C-terminal helix, producing a longer loop that remained largely disengaged from the TRP helix, maintaining only a distal contact (Glu703-Tyr409). While related remodeling occurred with mutation in the resting state, direct heating uniquely altered the S4-S5 linker/S5 helix junction (Fig. 4a), fusing them into a continuous helix.

**Figure 4.**
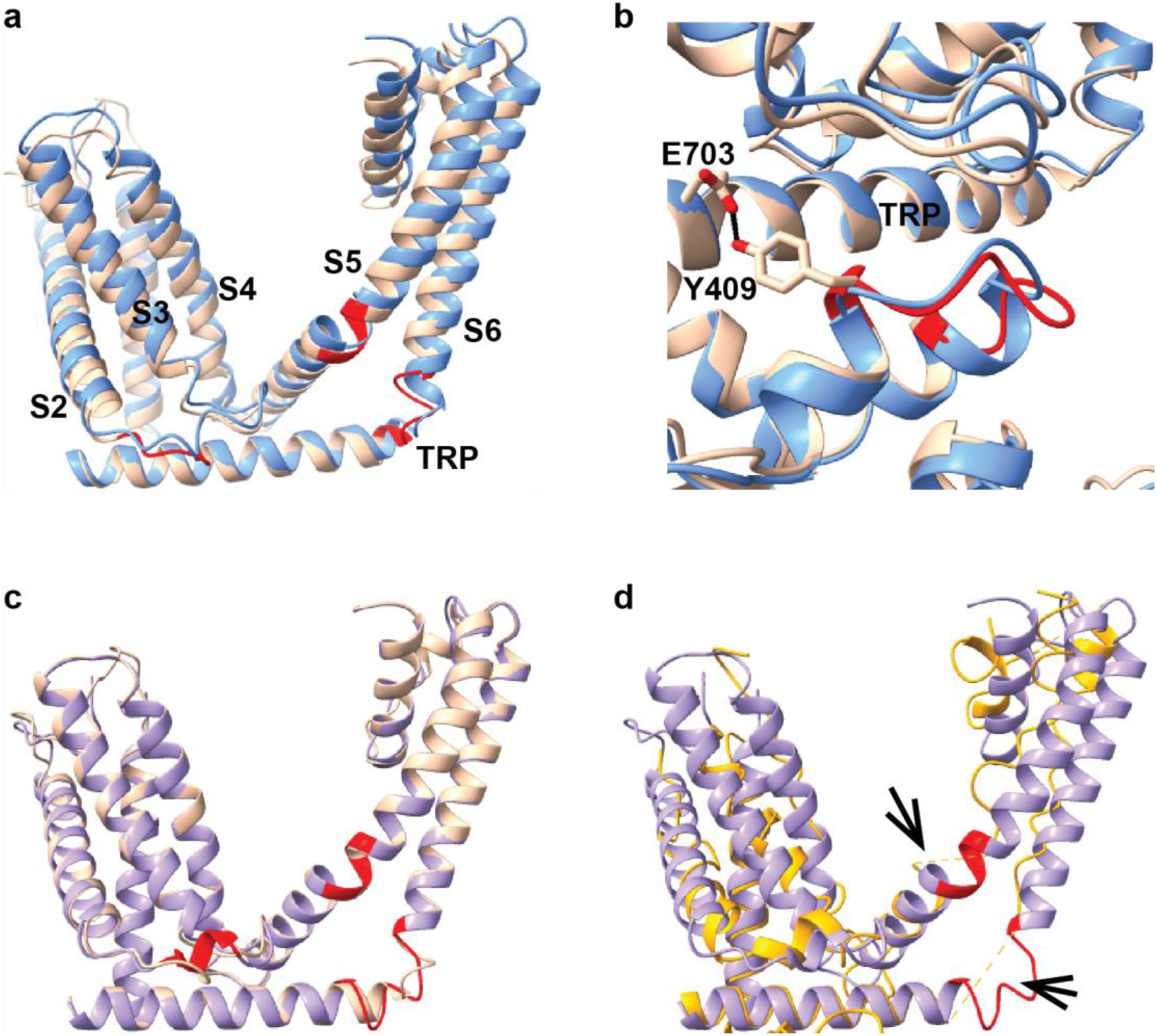
Progressive interfacial remodeling towards unfolding upon heating. **a-b)** Initial destabilization in the Vitrobot-heated structure (tan) overlaid with the resting TRPV3-412S structure (blue). (a) Overview of affected regions (red) in the tramsmembrane domain, including the S2-S3 loop, S6-TRP helix junction, and S4-S5 linker—S5 junction. These coincide with mutant-affected regions, with the additional transformation of the S4-S5 linker—S5 junction from a turn to a continuous fused helix. (b) Close-ups of TRP helix—proximal N-terminus interface, showing separation except for a single distal contact (Tyr409-Glu703). **c)** Progression of remodeling in DSC-heated (purple) versus Vitrobot-heated (tan) structures, highlighting helix formation in the S2-S3 loop and further unwinding at the S6-TRP elbow junction (red), while the fused S4-S5 linker with S5 helix remains. **d)** Mapping of remodeled interfaces to unfolding sites. Superimposition of the DSC-heated structure (purple) with the denatured structure (yellow) shows that the S4-S5 linker—S5 junction and the S6-TRP elbow junction in DSC-heated structure (purple) align with the regions where the pore detaches during unfolding.

In IM2, these changes were more advanced (Fig. 4c): the S2-S3 loop formed a defined helix (residues 511–514; Fig. 4c), consistent with prior heat-treated TRPV3 structures ^38^, and the N-terminal end of the TRP helix became disordered (Fig. 4c). This further extended the S6–TRP junction and increased its flexibility, which likely facilitated partial separation of the S6 bundle and contributed to the modest dilation of the lower gate at Met678 (Fig. S6-d). Importantly, distortions at the N- and C-terminal junctions of pore domain—the S4-S5 linker—S5 junction and the S6-TRP elbow—closely matched the rupture sites observed in the DSC-heated unfolded structure (Fig. 4d).

Globally, both heated structures exhibited annealing-like compaction (Fig. 5a-c), involving vertical shortening of ∼1 helix diameter, inward displacement of TMD towards the central pore, and counter-rotation of transmembrane helices relative to intracellular domains. This global contraction parallels that observed in the unfolded state (Fig. 1h,i) and in TRPV3 open-state structures ^37,38,54,61^. Subunit overlays revealed coordinated inward shifts of the S1–S4 bundle (Fig. S8-a), pore domain (Fig. S8-b), and TRP helix (Fig. S8-c), accompanied by an upward/inward rigid-body displacement of the ARD and C-terminal domains (CTD) by a ∼3–4 Å (Fig. S8-d). Importantly, individual domains maintained their internal structure, as confirmed by domain-specific superimpositions showing minimal changes within the S1-S4 bundle (Fig. 5d), pore domain (Fig. 5e), ankyrin repeats (Fig. 5f), proximal N-terminus (Fig. 5g), and C-terminus (Fig. 5h). Thus, the heat-induced compaction arose primarily from loops and interfacial remodeling between domains. Collectively, this pattern of changes—enhanced interfacial dynamics at the S6-TRP-MPD junctions coupled with loop remodeling driving global compaction while preserving domain rigidity—supports the view that temperature sensitivity depends on localized protein dynamics at key interfaces instead of wholesale domain rearrangements.

**Figure 5.**
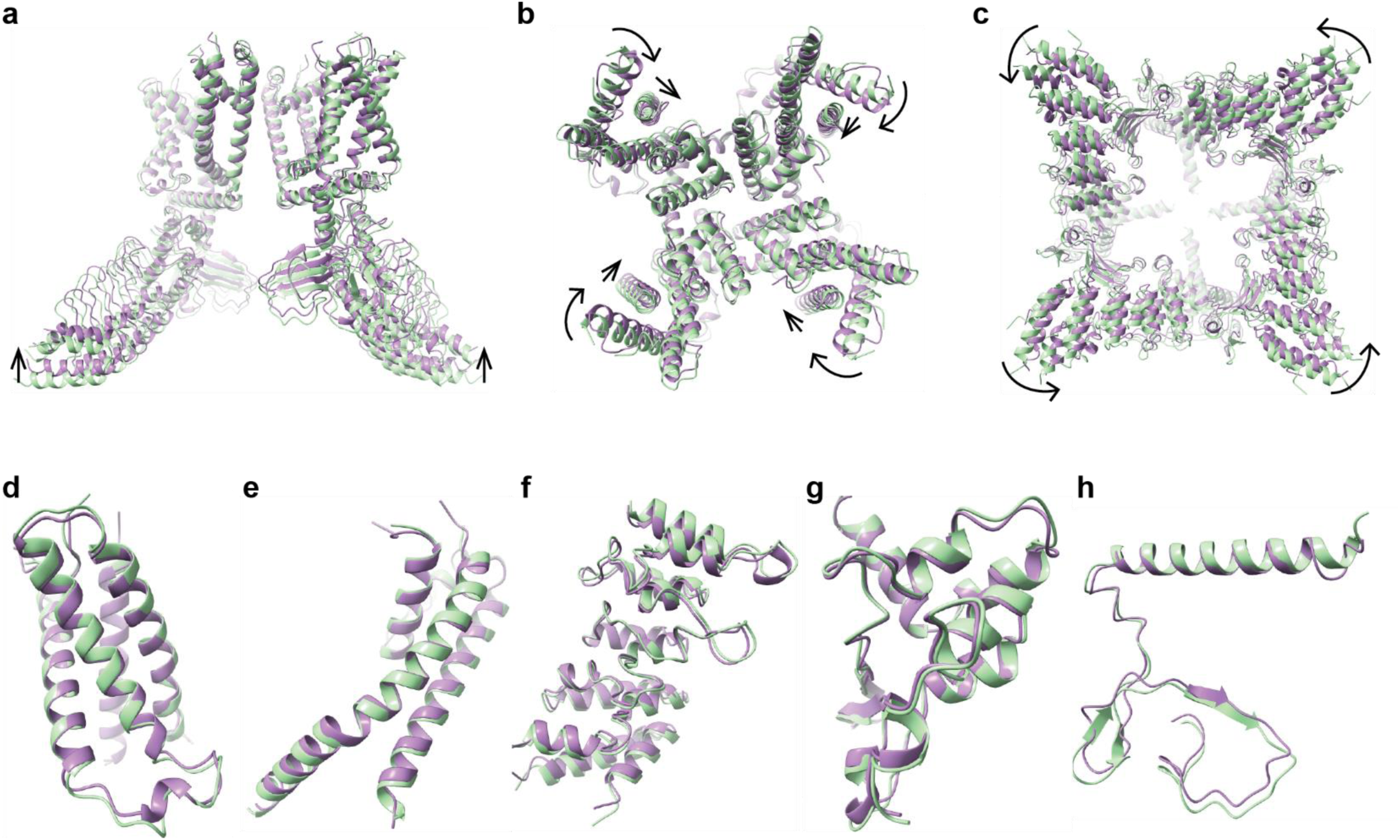
Global structural relaxation induced by thermal annealing. **a-c)** Superimposition of the DSC-heated structure (purple) onto the non-heated resting state (green) reveals coordinated global compaction and twisting movements: (a) side view shows a vertical contraction by ∼1 helix diameter (arrows); (b) top view of transmembrane domain shows compaction and a clockwise rotation (arrows); and (c) bottom view of intracellular ankyrin repeat domain (ARD) shows a counterclockwise rotation (arrows). **d-h)** Overlays of individual domains show minimal internal remodeling within the S1-S4 bundle (d), pore (e), ankyrin repeats (ARD1-4) (f), proximal N-terminus (g), and TRP helix (h), indicating that thermal annealing primarily modifies inter-domain orientations and linkages, while preserving the internal architecture of core domains.

### Thermolabile molecular latches in stepwise structural disassembly

The identification of four discrete states—resting, IM1, IM2, and denatured (DT)—suggests that TRPV3 undergoes progressive thermal transitions through structurally metastable intermediates rather than collapsing in a single catastrophic step. Tracking structural changes across this continuum revealed pivotal gating events and pinpointed thermolabile elements stabilizing the closed conformation. Central to this stability is a triad of electrostatic contacts in the membrane-proximal region that functions as a “latch”, linking the TRP helix, S6, and the S4–S5 linker to mechanically reinforces the lower gate and acts as an early barrier to thermal activation.

One key interaction occurs between Glu682 on TRP helix and Thr680 on adjacent TRP-S6 loop, forming an intersubunit ring of hydrogen bonds at the S6-TRP helix elbow (Fig. 6a). This network reinforces tetrameric assembly and secures the closed gate and remains in resting mutant (Fig. 6b) but becomes weakened in IM1 after heating and fully disrupted in IM2 (Fig. 6c). Its loss parallels progressive distortion of the S6–TRP helix loop (Fig. 4)—beginning with early destabilization in the mutant resting state, advancing to sequential unwinding of S6 and adjoining TRP helix coupled with loop elongation in IM1 and IM2, and culminating in DT with pore disassembly precisely at the TRP helix N-terminus—identifying this elbow junction as a thermally labile hotspot.

**Figure 6.**
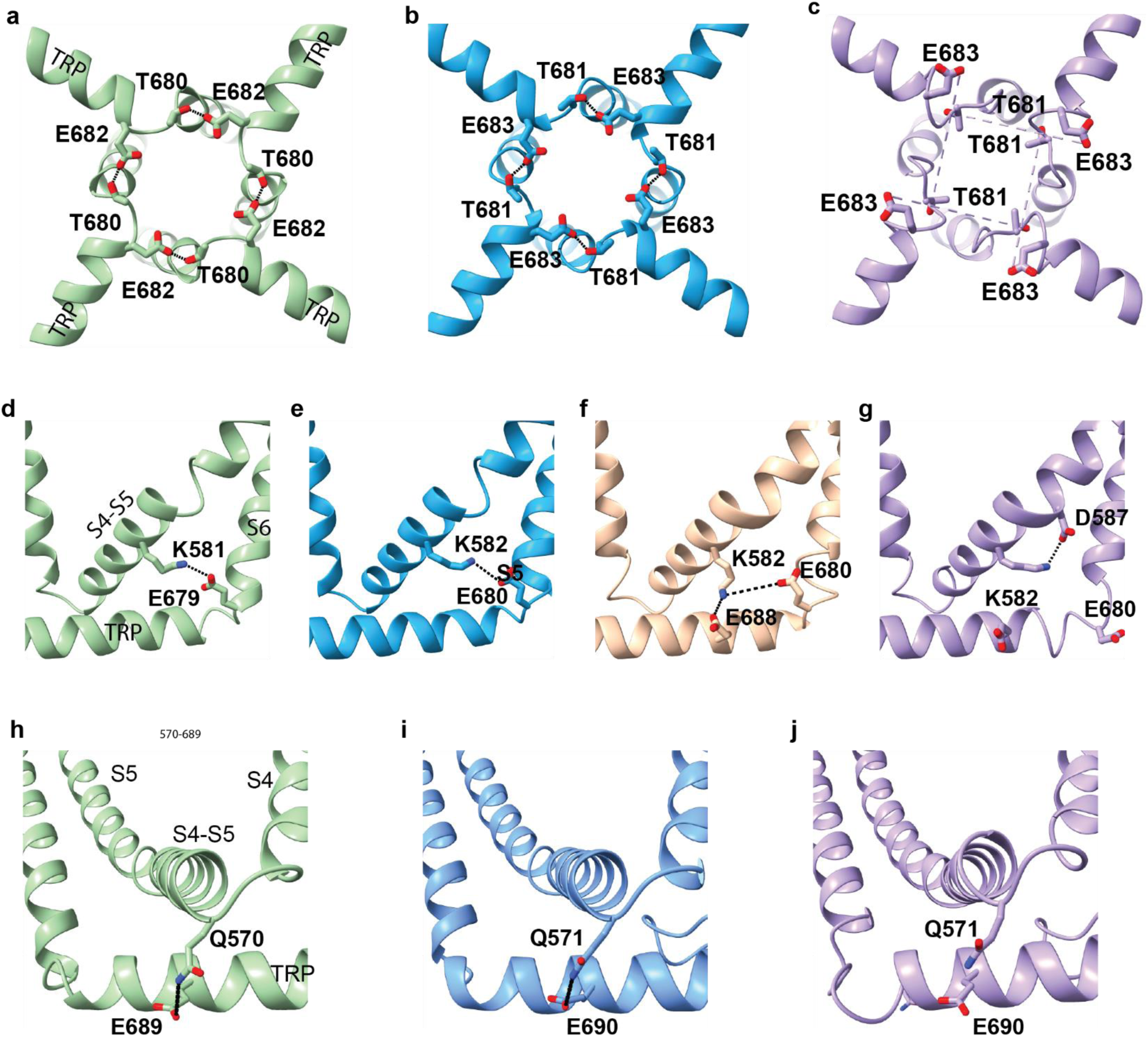
Stepwise destabilization of interfacial latches by heat. **a-c)** Intersubunit hydrogen bond Thr680—Glu682 linking S6 C-terminus to adjacent TRP helix N-terminus, forming a ring-like latch that stabilizes the tetramer. The contact lengthens progressively after heating: 2.6 Å (WT, a; mutant resting, b), 3.18 Å (Vitrobot), and is lost in DSC-heated structure (14.6 Å, c). **d-g)** Intrasubunit salt bridge Lys581 (S4-S5 linker)—Glu679 (S6 C-terminus), which stabilizes the S4-S5 linker connection to S6 in WT (3.5 Å, d), weakens in the mutant resting state (4.6 Å, e), and further lengthens after rapid heating (6.3 Å, f). Heating also rewires interactions: Lys581 forms a new salt bridge with Glu688 (TRP helix) after rapid heating (2.96 Å, f) and with Asp587 (S5) after DSC heating (3.53 Å, g). **h-j)** Intrasubunit hydrogen bond Gln570 (S4-S5 linker)—Glu689 (TRP helix) remains ∼3 Å in WT and mutant resting state (h, i) but ruptures in DSC-heated structure (>4 Å, j), indicating loss of the S4-S5 linker—TRP tether under prolonged thermal stress.

The N- and C-terminal ends of the pore domain, where rupture ultimately occurs, are further stabilized by a salt bridge between Glu679 (S6) and Lys581 (S4–S5 linker). In WT, these residues lie within interacting distance (3.6 Å; Fig. 6d). In the unheated mutant, the bridge persisted but at an increased distance (4.0 Å; Fig. 6e), suggesting weakened mechanical tethering. In IM1, brief heating broke the bridge as Lys582 reoriented toward Glu688 on TRP helix (Fig. 6f). This disruption coincided with helical fusion of S4-S5 linker and S5 (Fig. 4), signaling reduced mechanical constraint at this junction. By IM2, Lys582 rotated further to form a transient salt bridge with Asp587 on S5 (Fig. 6g), disengaging the S4-S5 linker from both the pore and TRP helix. This progressive loss of tethering frees the pore-lining S6 from peripheral stabilization, further weakening the closed state. Although linker fusion might suggest increased rigidity, the associated geometric strain likely exacerbate S5–S6 destabilization, predisposing the pore to denaturation.

A third contact, a hydrogen bond between Gln570 (S4–S5 linker)–Glu689 (TRP helix), additionally tethers the S4-S5 linker to the TRP helix and stabilizes the N-terminal end of the pore domain (Fig. 6h). While preserved in the unheated mutant (Fig. 6i) and IM1, this contact weakened markedly in IM2, with distance increasing from ∼3 Å to ∼4 Å (Fig. 6j), making another thermolabile hotspot susceptible to prolonged heating.

In parallel, the interaction between S2–S3 linker and proximal N-terminus (MPD), which normally supports TMD–cytosolic coupling and indirectly stabilizes the pore (Fig. 2f), was perturbed. Mutational disruption of this interface (Fig. 2g) impacted TRP helix and disordered the TRP–S6 junction (Fig. 2h). Upon heating, the S2–S3 linker progressively remodeled (Fig. 4a,c), transitioning from an incipient helix in IM1 to a defined short helix in IM2 ^38^, but without re-engaging the MPD. This trajectory effectively consolidated the decoupled state rather than restoring stabilizing contacts.

Collectively, these observations delineate a stepwise disassembly mechanism: Initial disruption of the intersubunit Glu683-Thr681 bond destabilizes the S6–TRP elbow, loosening the gate. Subsequent loss of Glu680–Lys582 salt bridge and Glu690–Gln571 interaction progressively decouples the S4–S5 linker from the TRP core, weakening transmembrane-cytosolic connectivity. Collapse of this stabilizing triad selectively undermines three key interfaces: the S4–S5 linker, S6–TRP junction, and S2–S3 linker/MPD– TRP hub. Concomitantly, a global annealing-like compaction, independent of heating rate, drives inward compression of peripheral S1–S4 bundles toward the central pore, while elongation of the S6–TRP helix loop expands the S6 bundle crossing. These coordinated local and global rearrangements likely intensify intersubunit tension and mechanical stress at pore-lining interfaces, promoting irreversible transition from a stable, closed conformation to mechanically compromised states. The conservation of this latch network in both WT and 412S channels suggests a shared disassembly pathway in WT, potentially accelerated by heat-sensitive MPD-TRP interactions.

## Discussion

Several conceptual models have been proposed to explain the extraordinary temperature sensitivity of thermoTRPs, including specialized sensor domains, nonpolar residue repartitioning, lipid modulation, and coupling to partial unfolding. Here, we provide the first direct structural evidence linking interfacial dynamics to steep temperature sensitivity. Rather than large-scale domain rearrangements, we observed only modest changes occurring at domain-domain interfaces, concentrated near the membrane-proximal region where the bundle crossing, TRP helix, proximal N-terminus, and intracellular faces of the S1-S4 bundle converge (Fig. 2e). In TRPV3, even selective disruption of these latches (Fig. 2g) markedly reduces wild-type temperature dependence, while heating destabilizes additional anchoring junctions (Fig. 4,6), ultimately leading to pore detachment and irreversible transitions (Fig. 1). This interfacial disassembly pathway reveals how local unfolding at the central pore couples structural destabilization to functional activation, highlighting intrinsic protein dynamics—particularly interdomain destabilization rather than changes within individual domains—as the principal driver of extreme temperature sensitivity

The disassembly pathway defined by thermolabile latching interfaces also unifies the broad, disparate residue distributions revealed by mutagenesis studies ^26-44^, which are scattered across the channel without clear convergence in location or chemistry—patterns incompatible with sensor-centric or heat capacity models ^20-22^. Although mutagenesis clearly implicates these sites in thermal sensitivity, their mechanistic roles have remained unclear. Our dynamics-driven framework now provides a unifying explanation: many reported sites, including the proximal N- and C-termini, TRP helix, pore domain, and S1–S4 bundle, map onto interfacial latches that stabilize the closed state (Fig. 2,4), while proximal N-terminus–ARD interactions further implicate distal ankyrin repeats (Fig. 2f,g; Fig. S4). Thus, both structural and mutagenesis evidence indicate that temperature sensitivity emerges not from a single specialized sensor domain but from distributed interdomain destabilization. Notably, although denaturation culminates at the central pore—possibly reflecting the instability of its inverted tepee architecture—structural-functional and calorimetric data show that extreme temperature dependence requires concerted destabilization of pore–interface networks ^26-28,49^. The pore may be intrinsically thermally sensitive; however, only cooperative interactions with surrounding domains generate the large activation energy barrier characteristic of thermoTRP gating.

Correlating structural changes with function is particularly challenging for thermoTRPs, whose activation is kinetically driven, involving hysteresis ^46^, sensitization ^48^, rundown ^50,52^, and denaturation ^49,50^, while cryo-EM captures only steady states. The interfacial latches we identified nonetheless align closely with with functional data, such as the helix-loop-helix interactions with the TRP helix that are critical for thermal sensitivity ^26^. Crucially, use of the sensitization-mimicking mutant allowed us to isolate changes specifically linked to temperature sensitivity. This mutant offered several advantages ^27^: its lower activation threshold minimized nonspecific effects from high temperatures stimulation; weakened interfacial coupling disrupted concerted activation ^49^, exposing localized unfolding events (Fig. 1,4); and, most importantly, it stabilized the transient sensitized closed state, the key intermediate where steep temperature dependence arises, whereas the final opening step is comparatively temperature-independent ^27,47^. By trapping this state without applying heat, standard cryo-EM could resolve structural correlates of temperature sensitivity while avoiding confounding transitions from pore opening or nonspecific unfolding. This likely explains why we observed localized interfacial changes, rather than the broader rearrangements reported seen in heated wild-type channels ^37,38,54^. Finally, our observation of a persistently α-helical S6 and lipid occupancy within an apparent fenestration tunnel– previously undescribed – suggests that cryo-EM may capture both intrinsic gating elements and preparation-dependent variations (Fig. 2i; Fig. S2 & S7).

In summary, our study provides a structural framework linking gating and unfolding in thermoTRP channels, directly capturing the destabilization events that underlie extreme temperature sensitivity. By mapping key molecular interactions and stepwise transitions, it reveals how modulators could selectively bypass or leverage latching interfaces to mitigate pathological thermal responses while preserving physiological sensing. More broadly, these findings highlight the central role of protein energetics in shaping channel gating and demonstrate that folding–unfolding transitions can be harnessed to achieve precise biological thermosensing.

## Materials and Methods

### Protein expression and purification

Mouse TRPV3 clone was kindly provided by Ardem Patapoutian (Scripps), and the BacMam expression vector was a gift of Erhu Cao (University of Utah). Protein expression and purification followed published protocols with modifications ^67^. Briefly, full-length channels were cloned into the BacMam vector ^68^ containing an N-terminal maltose binding protein (MBP) followed by a TEV protease cleavage site.

Recombinant baculovirus encoding the target genes were generated in SF9 cells using the Bac-to-Bac system (Invitrogen). Amplified P2 virus was used to infect HEK293 GnTI^-^ cells cultured in Freestyle293 media (Gibco) supplemented with 2.0 % FBS (Gibco) at 37 °C with constant shaking in a humidified 8% CO_2_ incubator. Cells were transduced at a density of ∼3.0 x 10^6^/ml.

Following infection, cultures were maintained at 37 °C for ∼24 hrs, then shifted to 30 °C, with 10 mM sodium butyrate added to enhance protein expression. After ∼48 h of infection, cells were harvested by centrifugation at 500 x g for 15 min and resuspended in hypertonic buffer (36.5 mM sucrose, 50 mM Tris, 2.0 mM TCEP, pH8.0) containing protease inhibitors (1.0 mM PMSF, 3.0 μg/ml aprotinin, 3.0 μg/ml leupeptin, 1.0 μg/ml pepstatin; all from Roche). Cell lysis was performed on ice for 30 – 45 min using a Parr bomb at 500 psi followed by nitrogen cavitation. Lysates were clarified by low speed centrifugation to remove intact cells and nuclei. Membranes were collected by ultracentrifugation at 100,000 x g for 1 hr and resuspended in wash buffer (WB: 200 mM NaCl, 2.0 mM TCEP, 10% glycerol, 20 mM HEPES, pH 8.0) with protease inhibitors.

Membranes were solubilized in 1% GDN (Anatrace) for 1 hr at 4 °C with gentle shaking, followed by centrifugation at 25,000 x g for 35 min to remove insoluble fractions. Supernatants were incubated with amylose resin (New England Biolabs) for 2 h at 4 °C, then transferred to a column, washed with WB containing 0.5mM GDN and 10 μg/ml soybean lipids (Avanti), and eluted with 20 mM maltose in the same buffer without glyceral. Purified protein homogeneity was assessed by size-exclusion chromatography (SEC) on a Superose 6 column equilibrated with WB containing appropriate detergents and lipids; samples were used immediately for reconstitution experiments.

### Nanodiscs reconstitution

Purified channel proteins were reconstituted into lipid nanodiscs for cryo-EM following established protocols ^69,70^, using membrane scaffold protein MSP2N2 for discs stabilization. The MSP2N2 plasmid, generously provided by Dr. Erhu Cao (University of Utah), was expressed in Escherichia coli BL21 cells and purified via nickel affinity chromatography ^71^. Eluded fractions were analyzed by SDS-PAGE, dialyzed extensively against 20 mM Tris/HCl, pH 7.4, 100mM NaCl, 0.5mM EDTA, and concentrated. Protein concentration was determined spectrophotometrically at 280 nm (ε = 21000 cm^−1^M^−1^).

For lipid preparation, soybean polar lipid extract (Avanti Polar Lipids) was dissolved in chloroform, dried under a nitrogen stream to form a thin film, and further desiccated under vacuum for 4-5 h to remove residual solvent. The lipid film was rehydrated overnight at 4 °C in reconstitution buffer (150 mM NaCl, 20 mM HEPES, 2.0 mM TCEP pH 7.4) at a final concentration of 7.0 mg/ml. Immediately before reconstitution, lipids were sonicated in a bath sonicator until optically clear.

Nanodisc assembly was initiated by combining TRPV protein with MSP2N2 and sonicated lipids at a molar ratio of 1:1:200 (TRPV monomer:MSP:lipid). The mixture was incubated on ice for 1 h. Detergent removal was then performed by sequential addition of three batches of pre-washed Bio-Beads SM2 (20 mg/mL; Bio-Rad) at 20 min intervals with gentle rotation at 4°C. TEV protease (40 μg per mg TRPV protein) was added in the final batch to cleave the MBP tag. The reaction was rotated overnight at 4°C to complete detergent removal and allow nanodisc self-assembly. The supernatant was separated from the Bio-Beads using a gel-loading pipette tip.

Assembled nanodisc-channel complexes were purified by size exclusion chromatography (SEC) on a Superose 6 Increase 10/300 GL column (GE Healthcare) equilibrated with SEC buffer (20 mM HEPES, 150 mM NaCl, 2 mM TCEP, pH 7.4). Fractions containing monodisperse complexes were pooled based on elution profiles and concentrated using Amicon Ultra centrifugal filters (Millipore) for downstream cryo-EM grid preparation.

### Cryo-EM sample preparation and data collection

Quantifoil 2.0/2.0 holey carbon grids (200-mesh) were used for standard preparations, while UltrAuFoil 1.2/1.3, Au-50 grids (300-mesh) were employed for in-chamber heating experiments. All grids were glow-discharged immediately before sample application using a PELCO easiGlow or EmiTech K100X (0.3 mBar, 15 mA, 25 s). Approximately 3 μL of protein (2-3 mg/mL) was applied to each grid. Vitrification was performed using a Vitrobot Mark IV (ThermoFisher Scientific) at >90% relative humidity, with a blot time of 4–8s under standard force settings, followed by plunge-freezing in liquid ethane. For non-heated samples, the chamber was maintained at 4°C. For rapid heating experiments, the chamber was pre-equilibrated to 42 °C, and grids were plunge-frozen within seconds of sample application to shorten thermal exposure. For slow heating experiments, samples were gradually heated in a differential scanning calorimeter (DSC) from ambient temperature (∼15 °C) to 42 °C at 1 °C/min, then centrifuged, and the supernatant was vitrified at room temperature.

Data were collected on 300kV Titan Krios microscopes (Thermo Fisher Scientific) equipped with Gatan K3 direct electron detectors at Brookhaven National Laboratory or Purdue University. Automated acquisition was performed using EPU software (Thermo Fisher Scientific) in super-resolution counting mode, with a nominal defocus range of 1.0–3.5 μm. Wild-type and mutant resting-state samples were imaged at a physical pixel size of 1.092 Å, using a dose of 1 e⁻/Å² per frame over 40 frames (total dose 40 e⁻/Å². Other samples were collected at 1.035 Å physical pixel size with 0.75 e⁻/Å² per frame across 67 frames (total dose 50 e⁻/Å²). Typical collection sessions lasted ∼24 h per sample and yielded 6,000– 9,000 micrographs.

### Single-particle data processing

Single-particle cryo-EM data were processed using cryoSPARC (Structura Biotechnology) following a similar workflow. Raw movie frames were aligned and dose-weighted using the Patch Motion Correction algorithm. Contrast transfer function (CTF) parameters were estimated from motion-corrected, summed micrographs via Patch CTF Estimation, and only micrographs exhibiting good CTF fits and resolution estimates were retained. Particles were automatically picked using cryoSPARC’s template-free blob picker. For initial processing, down-sampled particles (typically 4× Fourier binned) were extracted and underwent several rounds of reference-free 2D classification to remove junk particles and select coherent classes displaying clear TRPV channel features. The remaining particles were used for ab initio 3D reconstruction, and particles contributing to the best classes were re-extracted at full resolution. Iterative refinement in cryoSPARC included homogeneous refinement, non-uniform refinement, and further 3D classification to progressively improve map quality. To enhance conformational separation, particles were further exported to RELION 3.1 for focused 3D classification without alignment. Resulting subclasses were re-imported into cryoSPARC for additional rounds of refinement. Map resolutions were estimated from the two independently refined half-maps using the Fourier shell correlation (FSC) 0.143 cutoff. Local resolution variations were calculated using cryoSPARC or Phenix.

For wild-type channel, 8,114 micrographs were collected. Initial processing yielded 1,535,611 particles after 2D classification. Three ab initio 3D models were generated, and one (695,090 particles) was selected. This subset was exported to Relion 3.0 and subdivided into three 3D classes. The selected subsets were re-imported into cryoSparc for full-resolution re-extraction, additional 2D classification, and 3D refinement, producing a final map at 3.04 A resolution (C1 symmetry) from 234,297 particles.

For mutant resting-state dataset, 8,000 micrographs were collected. After 2D classification, 1,063,500 particles were selected. A single ab initio 3D model was generated and used for downstream refinement. Focused 3D classification in RELION 3.1 produced three distinct classes; particles from the best-resolved classes were re-imported into cryoSPARC for refinement with C4 symmetry, yielding a 2.72 Å resolution map from 288,168 particles.

For DSC-heated sample, 9,000 micrographs were collected, yielding 1,512,790 initially extracted particles (2x binning). One ab initio 3D model was generated in cryoSparc, and the particle set was exported to RELION for 3D classification into four classes. One class exhibited complete density and was re-imported into cryoSparc for full-resolution re-extraction, 2D cleanup, and 3D refinement, resulting in the IM2 map at 2.49 Å resolution from 387,682 particles. The remaining classes (1,109,224 particles), showing consistent partial densities, were re-extracted at full resolution. After 2D classification, 910,435 particles were used to generate a de novo 3D model, refined to 2.70 Å resolution (C4 symmetry). Further RELION 3D classification did not improve reconstructions. The IM2 structure represents ∼25% of total particles, while the denatured model accounts for ∼60%.

For Vitrobot-heated sample, 7,331 micrographs were collected. Particles were initially extracted with 2× binning. After 2D classification cleanup, 989,075 particles were retained to generate a de novo 3D model. The dataset was then exported to RELION for 3D classification under C1 symmetry (five classes). The best-resolved subclass (208,236 particles) was re-imported into cryoSPARC for full-resolution re-extraction and homogeneous/non-uniform refinement, yielding a final reconstruction at 2.84 Å resolution.

### Model building

Initial model placement was performed in UCSF Chimera using previously determined TRPV3 structures (e.g., PDB ID: 6DVW) as starting references. Models were manually docked into the cryo-EM density maps, followed by rigid-body refinement using the Fit in Map tool to optimize model-to-density correlation. The channel was segmented into discrete domains—N-terminus, S1–S4 bundle, pore, and C-terminus—and each domain was individually refined using cryo-EM real-space refinement in Phenix. After domain-wise fitting, the complete model was reassembled and subjected to iterative manual real-space refinement in Coot, including adjustments to side chains, backbone geometry, and secondary structure elements. Site-directed mutations were introduced in Coot via real-space residue replacement, followed by rotamer optimization and steric clash minimization. For flexible regions, multiple conformations were modeled where density permitted. In particular, the HLH loop of TRPV3-412S (residues Asn410–Ile414) exhibited density consistent with potential conformations. The main and second conformations allowed assignment of side chains at a contour level of 3–4 RMSD, while the third conformation permitted tracing of the main chain only. The quality of the final models was evaluated using MolProbility, and pore radius profiles and constriction sites were analyzed with HOLE2, using a probe radius of 8.0 Å. Molecular graphics and figures, including atomic models and density overlays, were generated in ChimeraX (UCSF).

## Acknowledgement

This work was supported by the National Institutes of Health (NIH) grant R01 GM132762. The authors are grateful to Dr. Qiu-Xing Jiang for valuable experimental assistance.

## Legends for Supplemental Figures are included inline beneath each figure

**Figure S1.**
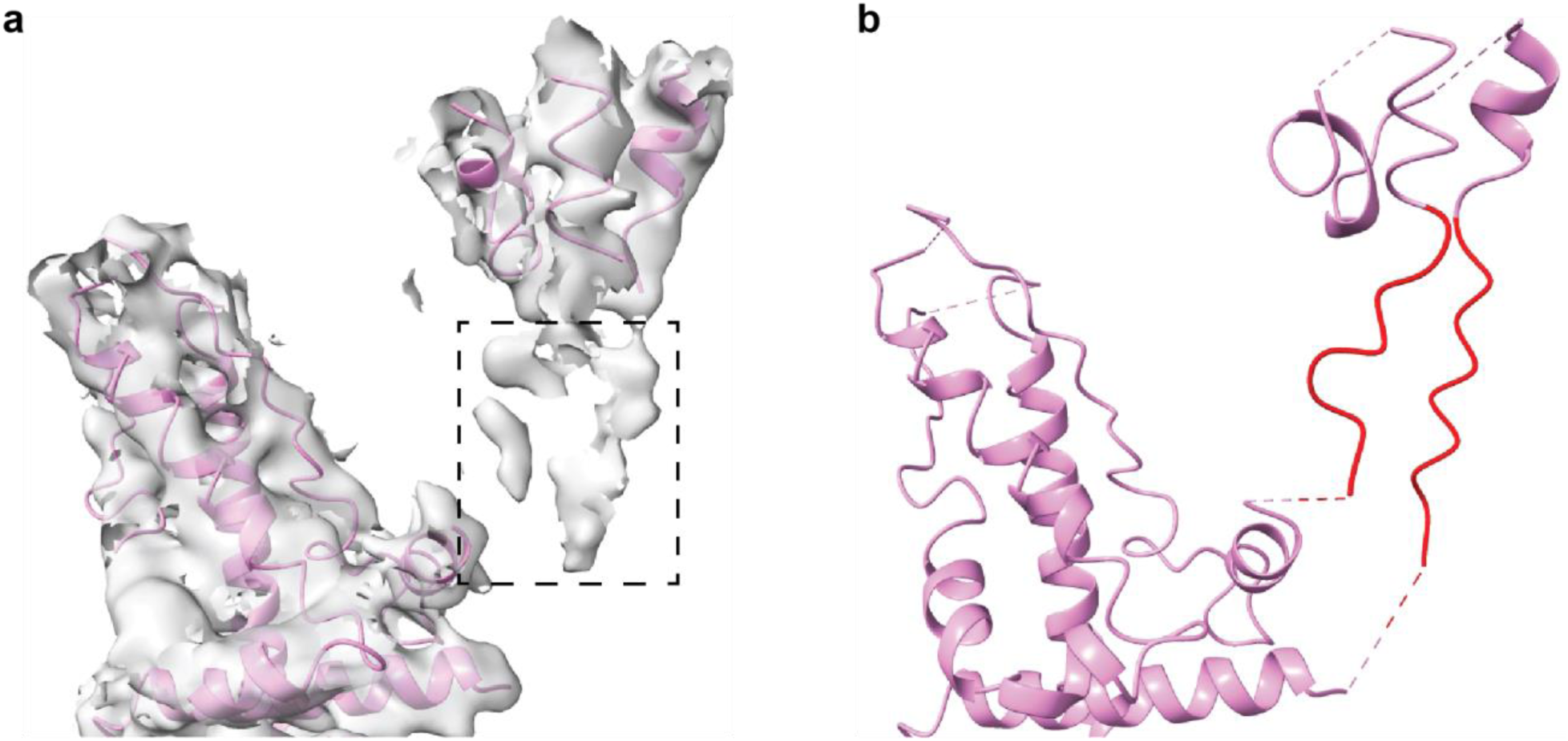
Partial backbone modeling of the pore. **a)** 3D density map of the pore region contoured at lower threshold, revealing weak, partial density along the S5 and S6 helices (boxed). **b)** Backbone model fitted to the extra density, partially accounting for residues 602-595 in S5 and 661-668 in S6 (red). The density decreases toward the intracellular side, indicating unfolding is most pronounced in the lower pore domain.

**Figure S2.**
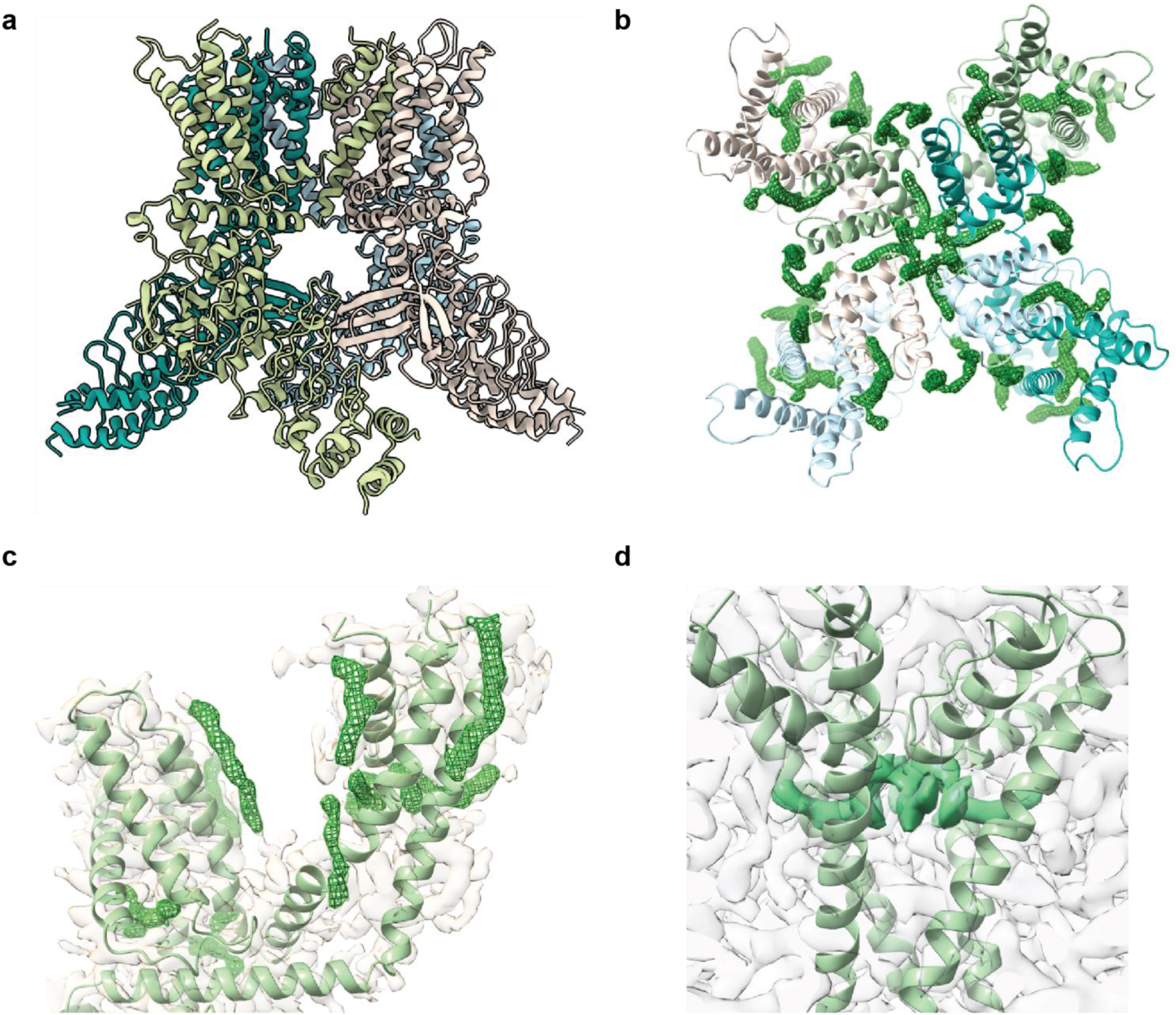
Wild-type TRPV3 structure in nanodiscs. **a)** Atomic model of wild-type TRPV3 reconstituted in lipid nanodiscs (overall 2.9 Å resolution), with individual subunits color-coded. **b)** Top view highlighting lipid-like densities associated with both peripheral S1-S4 bundles and central pore domain. Peripheral densities occupy (i) an outer-leaflet-facing crevice between the extracellular half of S4 and an adjacent subunit pore domain, and (ii) an inner-leaflet-facing pocket between the intracellular side of S1-S4 bundle and TRP helix. Central densities form a ring-shaped belt beneath the pore helix, with “finger-like” extensions penetrating lateral fenestrations at intersubunit interfaces between S6 and adjacent S5/S6 helices. **c)** Side view showing the spatial distribution of lipid like densities across extracellular, lateral and intracellular membrane leaflets. **d)** Magnified side view of the ring-shaped density, emphasizing its projections into lateral fenestrations.

**Figure S3.**
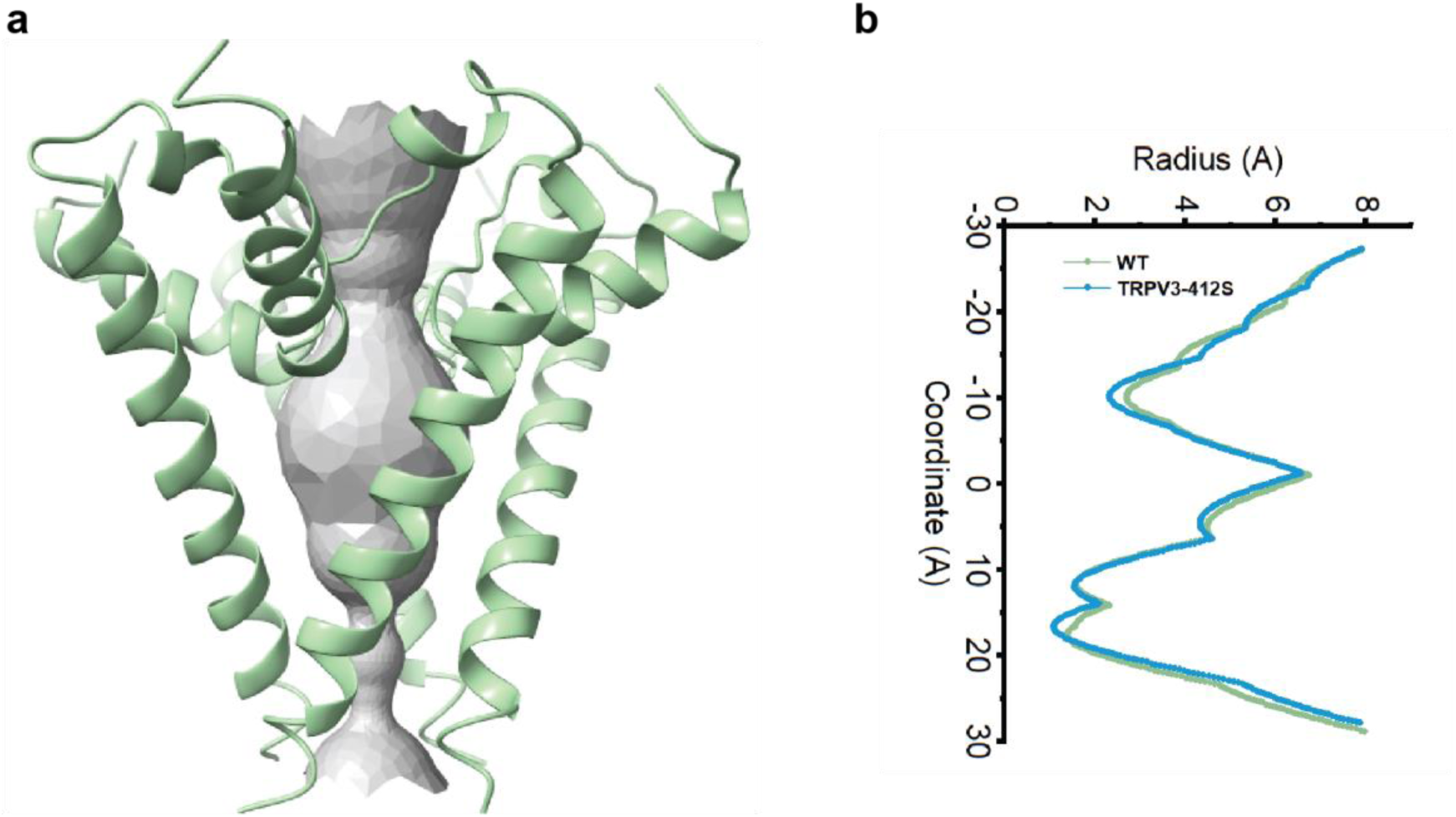
Pore radius profile of wild-type TRPV3. **a)** Pore geometry of resting wild-type (WT) TRPV3 visualized along the central channel axis, with pore-lining surface (grey) computed using the HOLE2 program, estimating the solvent-accessible radius at each axial position without intersecting the van der Waals surfaces of lining atoms. **b)** Superimposed pore radius profiles for wild-type (green) and TRPV3-412S (blue) and in resting states, both displaying comparable constrictions at the selectivity filter and lower gate (S6 bundle crossing), consistent with a closed pore conformation.

**Figure S4.**
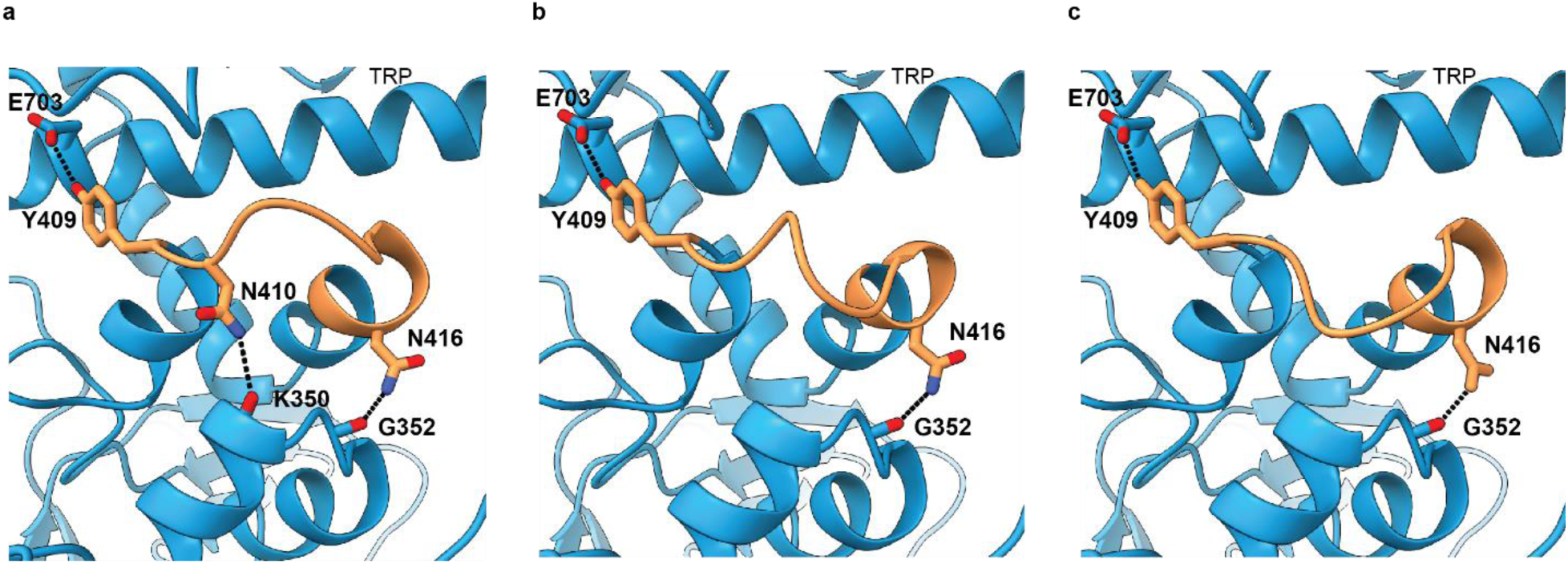
Conformational variability and weakened coupling between TRP helix and N-terminus in TRPV3-412S. Focused cryo-EM density modeling resolved three alternate conformations (a-c) of the loop (residues 408–414, orange) within the helix-turn-helix motif (HLH; residues 403–422) of proximal N-terminus, which lies between TRP helix and ankyrin repeat domain (ARD) and bridges intracellular N terminus to transmembrane domain (TMD). In wild-type (WT), this loop forms two contacts with TRP helix and two with ARD (Fig. 2e), and one with S2-S3 loop, whereas TRPV3–412S retains only a single distal TRP helix contact (Tyr408-Glu703) in all conformers. Of the two ARD-loop contacts (Lys350–Asn410 and Gly352–Asn416), both are preserved in Conformer 1 but in Conformers 2 and 3 only Gly352–Asn416 remains while Lys350–Asn410 is lost, further weakening reducing N-terminal coupling.

**Figure S5.**
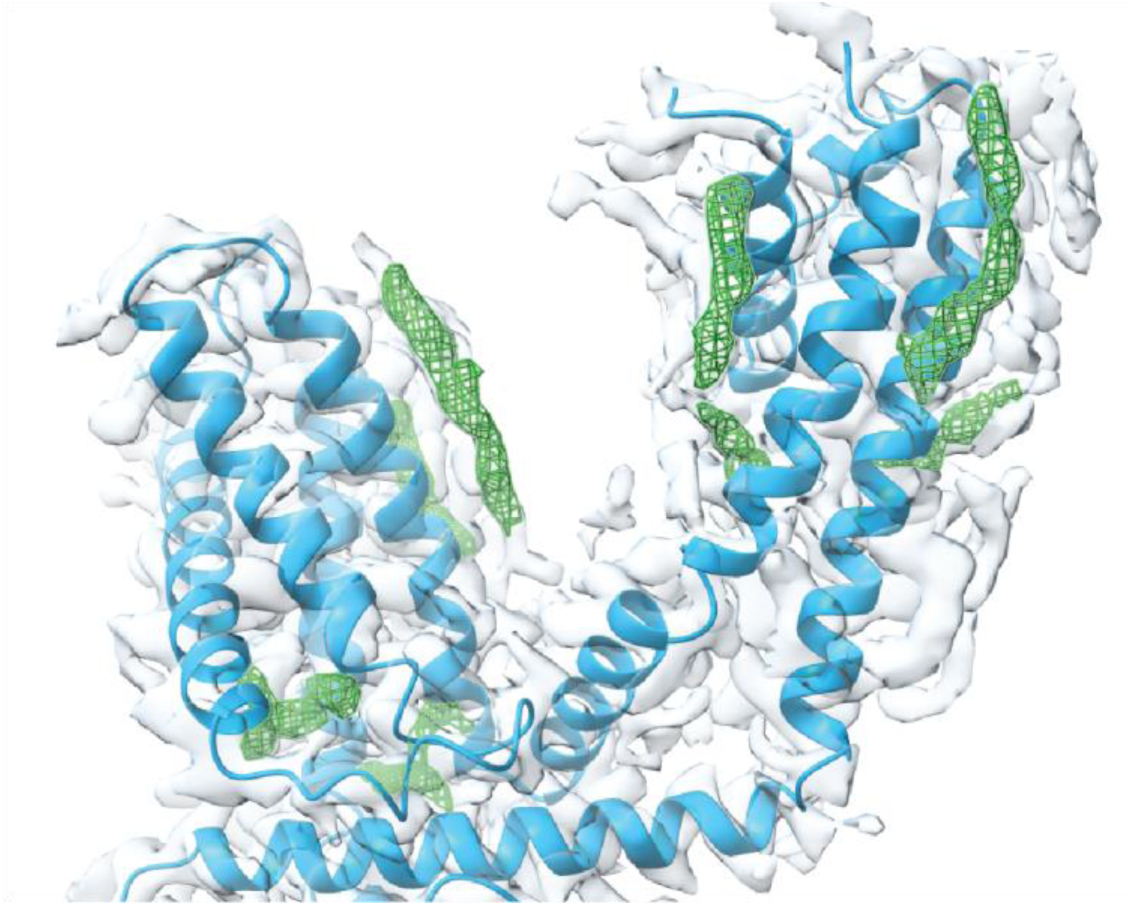
Lipid-like densities in TRPV3-412S. Side view showing their distribution (green) in the pore region, S1-S4 bundle, and vanilloid binding cavity.

**Figure S6.**
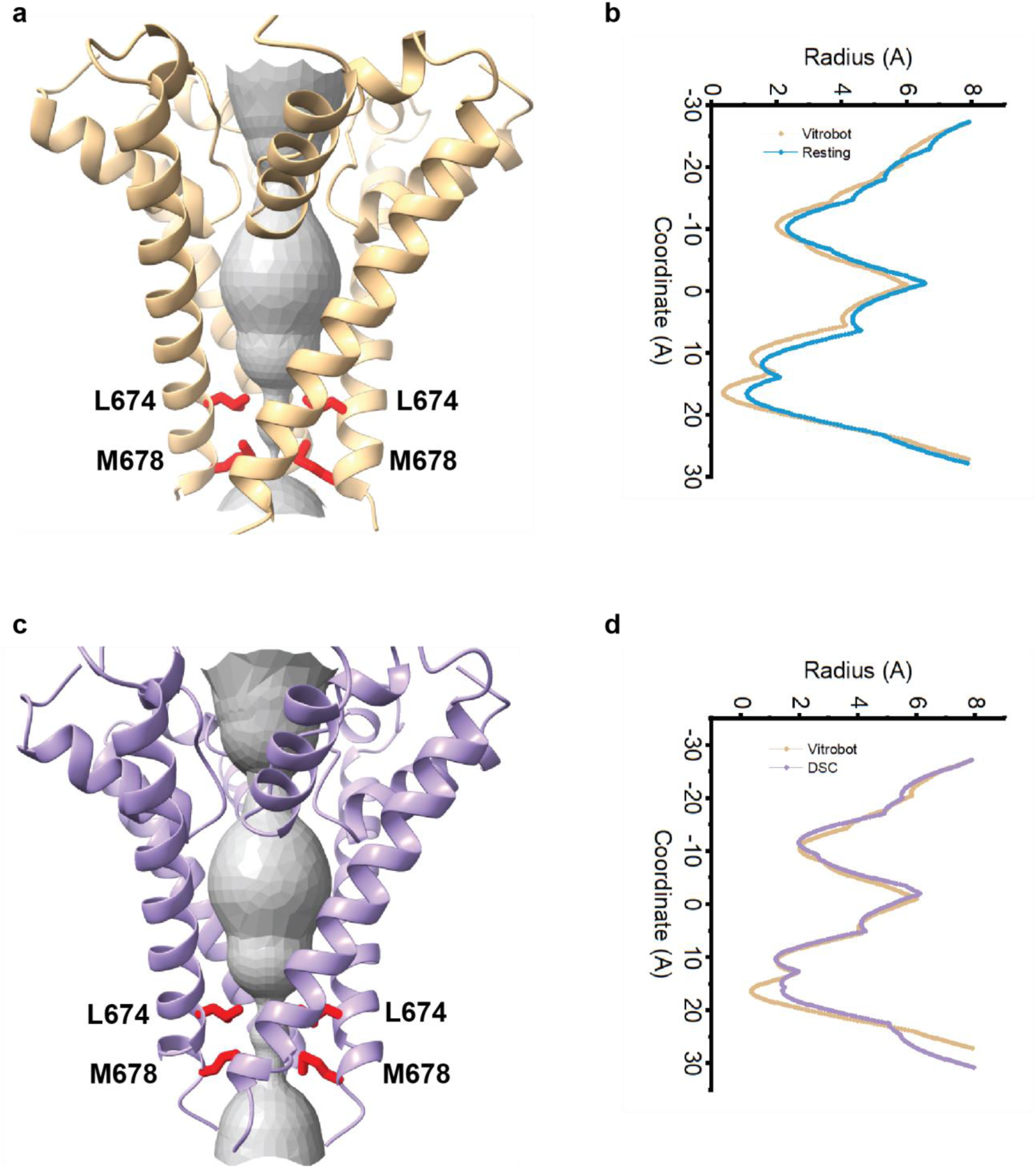
Pore radius profiles of heated TRPV3-412S structures. **a)** Space-filling model of the pore in Vitrobot-heated sample along the channel axis. **b)** Corresponding pore radius profile (tan) overlaid on non-heated control (blue). **C)** Space-filling view of the pore in DSC-heated sample. **d)** Pore radius profile (purple) superimposed on Vitrobot-heated profile (tan).

**Figure S7.**
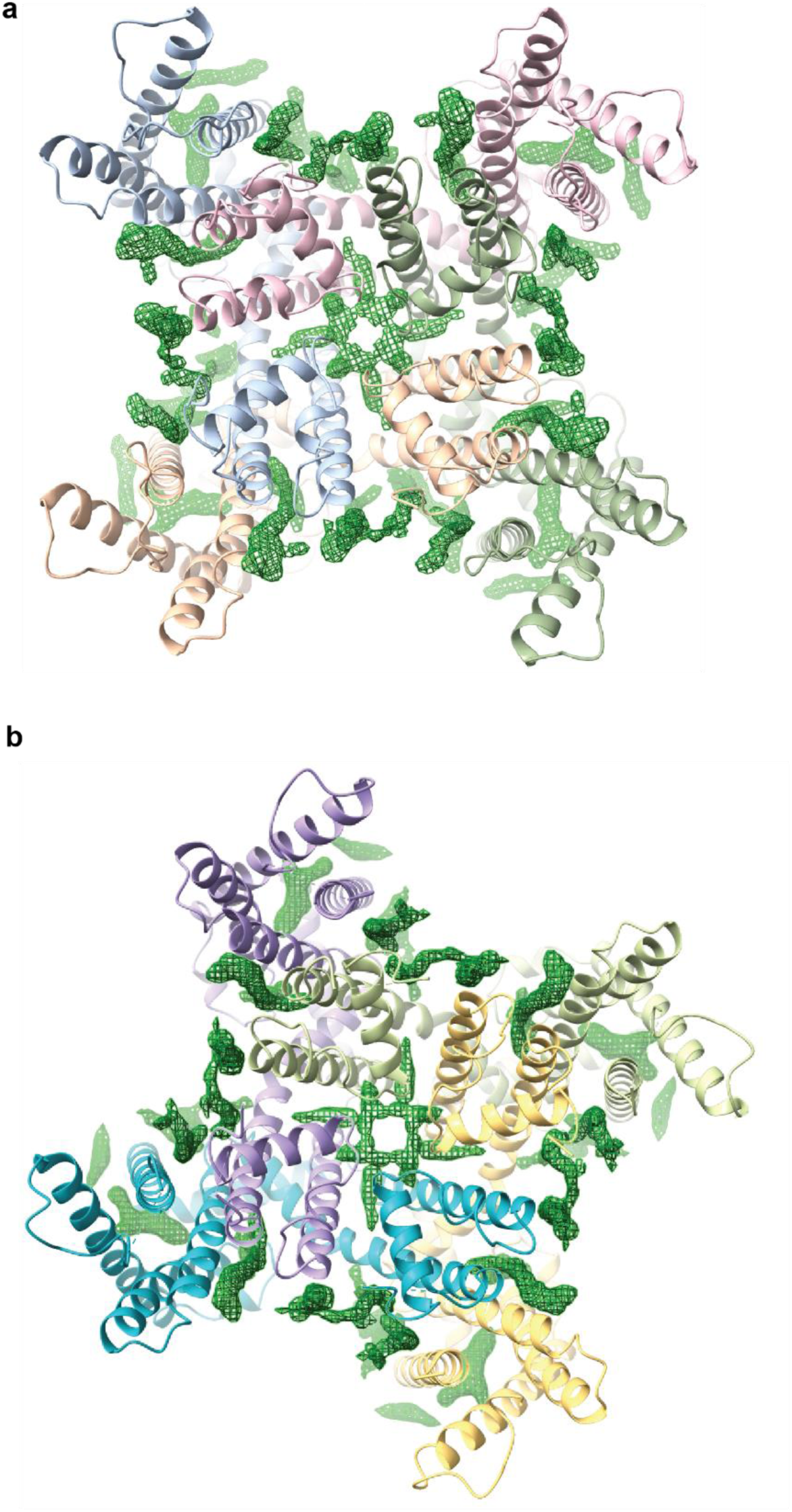
Lipid-like densities in heat-activated TRPV3-412S intermediates. **a)** Vitrobot-heated intermediate (IM1). **b)** DSC-heated intermediate (IM2). Both cryo-EM reconstructions exhibit non-protein densities (green) consistent with bound lipids at peripheral and central pore domains, analogous to those observed in wild-type (Fig. S2) and in non-heated mutant resting states (Fig.2c, Fig.S5), suggesting that heating maintains the overall lipid environment at these positions without gross rearrangement.

**Figure S8.**
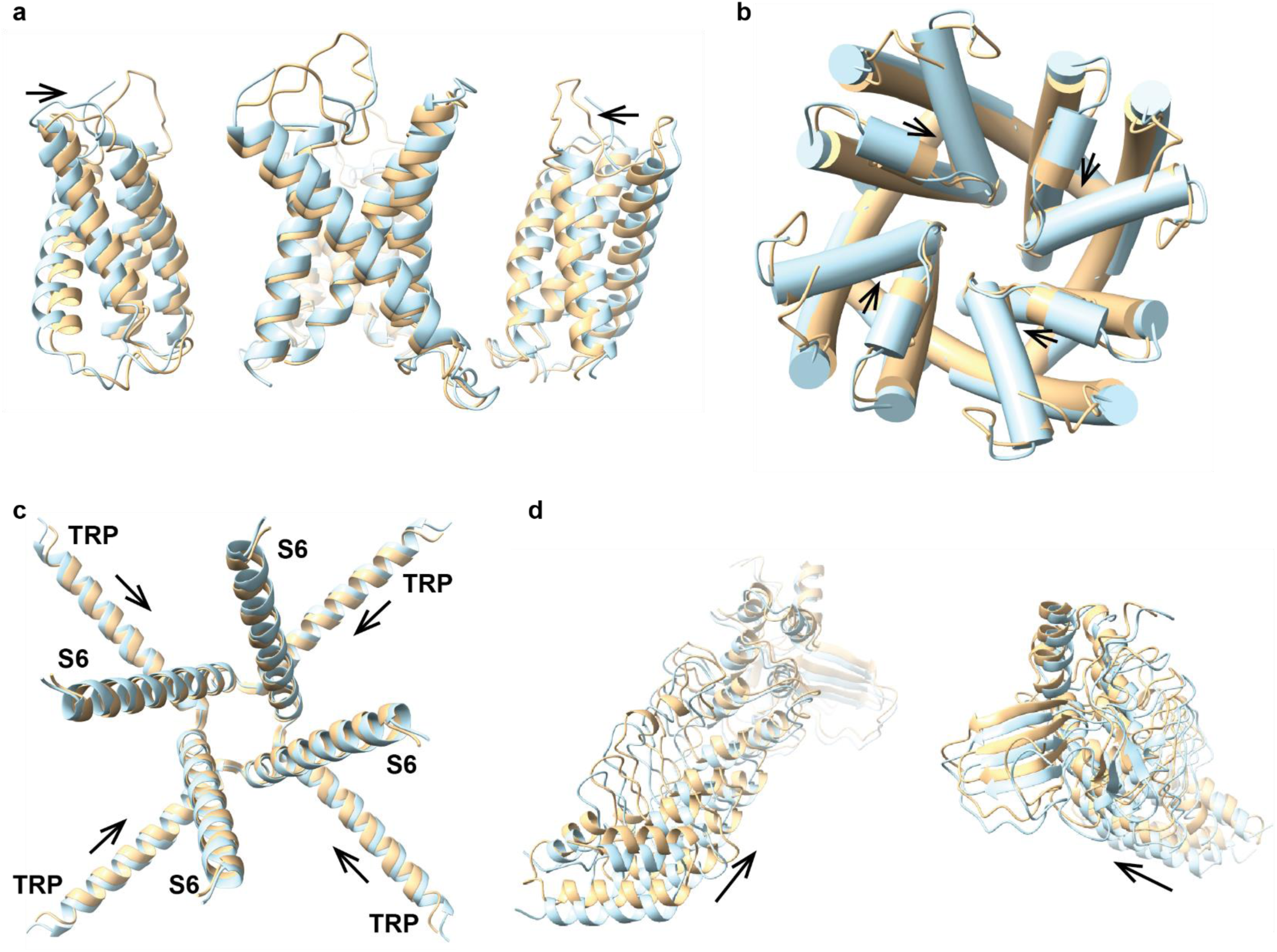
Global compaction during heat activation. **a)** Overlay of S1-S4 bundles from heat-activated intermediate (IM1; tan) and unheated state (blue), showing consistent inward displacement toward the central pore axis across subunits. **b)** Similar contraction of pore domain (S5-S6), most evident around pore helix (arrows). **c)** Top view of TRP helix and S6, illustrating an inward shift of TRP helix after heating (arrows). **d)** Side view of ankyrin repeat domain (ARD) and C-terminal β-sheets, showing coordinated upward and inward movement (arrows). All structures were aligned on their cryo-EM density maps prior to model docking. IM2 closely superposes with IM1 (Fig. 3g,h), confirming this global compaction, involving coordinated transmembrane and intracellular motions while preserving intradomain integrity, is conserved during heat activation.

## Notes

### Competing Interest Statement

The authors have declared no competing interest.

